# Creating basis for introducing NIPT in the Estonian public health setting

**DOI:** 10.1101/431924

**Authors:** Olga Žilina, Kadri Rekker, Lauris Kaplinski, Martin Sauk, Priit Paluoja, Hindrek Teder, Eva-Liina Ustav, Neeme Tõnisson, Konstantin Ridnõi, Priit Palta, Kaarel Krjutškov, Ants Kurg, Andres Salumets

## Abstract

**Objective:** The study aimed to validate a whole-genome sequencing-based NIPT method and our newly developed NIPTmer analysis software with the potential to integrate the pipeline into prenatal clinical care in Estonia.

**Method:** In total, 447 maternal blood samples were included to the study. Analysis pipeline involved whole-genome library preparation and massively parallel sequencing on Illumina NextSeq 500. Aneuploidy status was determined with NIPTmer software, which is based on counting pre-defined per-chromosome sets of unique k-mers from raw sequencing data. To estimate fetal fraction (FF) from total cell-free DNA SeqFF was implemented.

**Results:** NIPTmer software allowed to identify correctly all samples of non-mosaic T21 (15/15), T18 (9/9) and T13 (4/4) cases. However, one mosaic T18 remained undetected. Six false positive results were observed, including three for T18 (specificity 99.3%) and three for T13 (specificity 99.3%). FF < 4% (2.8-3.99%) was estimated in eight samples, including two samples with T13 and T18. Despite low FF, these two samples were determined as aneuploid with NIPTmer software.

**Conclusion:** Our NIPT analysis pipeline proved to perform efficiently in detecting common fetal aneuploidies T21, T18 and T13 and is feasible for implementation into clinical service in Estonia.

## Introduction

Aneuploidies are a major cause of perinatal morbidity and mortality. In many countries, including Estonia, prenatal testing for fetal aneuploidies relies on analysis of maternal serum biomarkers combined with ultrasound examination (combined testing, CT) after which women who are deemed to be at high risk are offered an invasive confirmatory test. The main issue associated with this scheme is relatively high false-positive (FP) rate of CT due to which around 90% of the women, eligible for the following invasive testing, carry a healthy baby. At the same time, invasive methods – amniocentesis, chorionic villus sampling (CVS) and umbilical cord puncture – carry a risk of 0.1-0.2% to cause an abortion, even if the procedure is carried out by an experienced specialist (1). The number of unnecessary invasive testing can be reduced significantly if more precise primary screening methods are available. The breakthrough in the field came in 2008 with introduction of non-invasive prenatal testing (NIPT), when it was demonstrated that fetal aneuploidies can be detected by analysis of cell-free DNA (cfDNA) circulating in maternal blood and representing a mixture of maternal and fetal cfDNA (2, 3). The majority of cell-free fetal DNA (cffDNA) originates from the outer cytotrophoblast and therefore, in fact, represents the genetic makeup of a placenta (4). CffDNA is detectable in maternal plasma from as early as 5 weeks gestation and depending on gestational age constitutes 10-40% of the total plasma cfDNA (5–8). Since 2011, NIPT is routinely used for the detection of common fetal trisomies, such as trisomy 21 (T21), 18 (T18) and 13 (T13), demonstrating high sensitivity and specificity up to >99% (9, 10). Presently NIPT testing is growing constantly worldwide as it is recognized as a very sensitive and precise screening test that is not more hazardous for a pregnant woman than a simple blood draw procedure.

Given that NIPT offers a more accurate selection of high-risk pregnancies and enables to avoid a large number of unnecessary invasive procedures, we aimed to validate a whole-genome sequencing-based NIPT method and our newly developed NIPT analysis software (11) with the potential to integrate it into the prenatal clinical care in Estonia.

## Methods

### Samples and study design

A total of 431 samples from pregnant women were collected at Tartu University Hospital (Tartu, Estonia) and at East-Tallinn Central Hospital (Tallinn, Estonia) during 2015 to 2018. Gestational age ≥ 10 weeks was the mandatory requirement for enrolment. The study was approved by Research Ethics Committee of the University of Tartu (#246/T-21) and written informed consent was obtained from all participants. All experiments were performed in accordance with relevant European guidelines and regulations.

The cohort included cases with both high (referred as “high-risk” pregnancies; n=259) and low (referred as “general” population; n=172) risk of aneuploidy. High risk factors included increased trisomy risk of >1/300 as suggested by typical screening procedures, family history of aneuploidy, ultrasound abnormality, and maternal age over 36 years. All women from the “high-risk” group underwent invasive prenatal diagnostic procedure (amniocentesis or chorionic villus sampling followed by karyotyping, FISH or chromosomal microarray analysis, CMA). Pregnant women from “general” population revealed negative result on the first trimester screening; neither they had any detectable fetal abnormalities on ultrasound examination. In this case, the general health status of a baby was estimated on birth by neonatologist. The mean age of participants was 33 years (median 32, range 16-47) and gestational ages were in range of 10 to 40 weeks, with a median of 16 (11-21 weeks with a median of 15 for “high-risk” group; 10-40 weeks with a median of 25 for “general” population). General characteristics of the study cohort are summarized in Supplementary Table I.

Samples from “general” population were analysed by massive-parallel sequencing (MPS) to generate preliminary reference dataset. This was initially used to analyse the data obtained from the “high-risk” group enriched with aneuploid samples. The reference dataset was updated constantly by including euploid samples from “high-risk” group, which makes further analysis more precise.

In addition, 16 clinical cfDNA samples from the Center for Human Genetics, UZ Leuven, Belgium enriched for aneuploid cases were blindly analysed using the same method. The results were then compared with KU Leuven laboratory results.

The study design was a blind testing where karyotypes of the fetuses were not available in advance to the lab personnel. No results were returned to the participants of the study.

### Sample processing and sequencing

Samples (n=447) were processed according to previously published guidelines with minor modifications (12). Briefly, the procedure was as follows: peripheral blood samples were collected in cell-free DNA BCT tubes (Streck, USA) and plasma was separated within 72 hours by standard dual centrifugation. cfDNA was extracted from 4 to 5 ml plasma using QIAamp Circulating Nucleic Acid Kit (Qiagen, Germany). Whole-genome libraries were prepared using TruSeq ChIP Library Preparation kit (Illumina Inc., San Diego, CA, USA) as described in Bayindir et al., with 12 cycles for the final PCR enrichment step (12). Following quantification, equal amounts of 24 libraries were pooled and the quality of the pool was assessed on Agilent 2200 TapeStation (Agilent Technologies, Santa Clara, CA, USA).

Sequencing was performed on the NextSeq 500 platform (Illumina Inc.) with an average coverage of 0.32x (minimum 0.08, maximum 0.42) producing 85 bp single-end reads.

### Fetal DNA fraction calculation

Sequencing read data was mapped with bowtie 2.3.4.1 (13) against GRCh37 pre-built bowtie index file with pre-set option of ‘very-sensitive’ and the maximum fragment length of 500 for valid paired-end alignments (14). Mapped data was filtered by mapping quality of 30, after which SeqFF (14) with minor modifications (to optimise the file handling in SeqFF calculation workflow) was used to calculate fetal DNA fraction for all samples.

### Detection of fetal trisomies

The data was analysed with our novel NIPTmer software package, which is based on counting pre-defined per-chromosome sets of unique k-mers from raw sequencing data and applying linear regression model on the counts, thus avoiding time-consuming read-mapping step. NIPTmer calls aneuploidies of the chromosome of interest based on the z-score (difference between expected and observed values, normalized to the standard deviation of expected values) of the detected fraction of k-mers compared to the expected fraction predicted by model. Thus z-score 1 corresponds to about 15% probability of given observation coming from normal karyotype group, z-score 2 about 2% and so on. The details of the analysis as well as the validation of the software can be found elsewhere (11).

## Results

A total of 447 samples were analysed with an established NIPT pipeline. Fetal DNA fraction estimates by SeqFF were between 3–49% (with median of 11%), and average FF was lower in the “high-risk” group compared to the “general” population group (10% (range 3-29%) vs 17% (range 5-49%), respectively), which can be explained by a wider gestational age range of the latter.

Z-score threshold for common trisomies (trisomy 21 (T21), 18 (T18) and 13 (T13)) was set at 3.0, which allowed to identify correctly all positive samples for non-mosaic T21 (15/15), T18 (9/9) and T13 (4/4). However, one mosaic T18 remained undetected (z-score 2.43). FISH of chorionic villus revealed the presence of T18 in ~20% of the cells, while karyotyping of the same sample was not able to establish the percentage of mosaicism because of the small amount of the cells analysed. Six FP results were observed, including three for T18 (specificity 99.3%) and three for T13 (specificity 99.3%). Four FPs were detected among “general” population pregnancies.

Invasive test detected one mosaic trisomy 8 (karyotype 47,XY,+8(11)/46,XY(29)), but there was no evidence of it with NIPT. No other autosomal trisomies, neither chromosomal deletions nor duplications were observed by amniocentesis/CVS. The results are summarized in Table 1.

**Table 1.** NIPT results for common aneuploidies (T21, T18, T13) and follow-up data of the study population (n=447).

|  | Estonian cohort |  | Belgium cohort | Total | FP | FN |
| --- | --- | --- | --- | --- | --- | --- |
|  | High-risk group | General population | Samples enriched for aneuploidies |  |  |  |
| Number of samples | 259 | 172 | 16 | 447 |  |  |
| T21 |  |  |  |  |  |  |
| NIPT | 13 | 0 | 2 | 15 | 0 | 0 |
| Invasive prenatal diagnosis | 13 | - | 2 | 15 |  |  |
| Neonatal-follow-up | - | 0 | - |  |  |  |
| T18 |  |  |  |  |  |  |
| NIPT | 8 | 1 | 3 | 12 | 3 | 1* |
| Invasive prenatal diagnosis | 7 | - | 3 | 10 |  |  |
| Neonatal-follow-up | - | 0 | - |  |  |  |
| T13 |  |  |  |  |  |  |
| NIPT | 1 | 3 | 3 | 7 | 3 | 0 |
| Invasive prenatal diagnosis | 1 | - | 3 | 4 |  |  |
| Neonatal-follow-up | - | 0 | - |  |  |  |

FP - false-positive results; FN - false-negative results
\* mosaic T18 case (20%)

Fetal sex (male or female) was determined correctly for all samples where the information was available (397/397; Supplementary Table 1). Five cases with sex chromosome aneuploidies (SCA) from “high-risk” group were identified by invasive testing of which four monosomy X (45, X0; Turner syndrome) cases were determined by NIPT (z-scores for X chromosome between −8.1 to −16.4), however one 47, XXX karyotype remained undetected (z-score −3.3).

For eight samples the proportion of fetal cfDNA in maternal plasma remained below 4% (2.8-3.99%), which is a widely accepted minimum that allows reliable detection of common trisomies (15–17). Six of these samples were from “high-risk” pregnancies and two were from Belgium cohort. Despite low FF in two aneuploid samples from Belgium, we were able to correctly identify T13 and T18 cases. All the rest six cases with FF below 4% from “high-risk” Estonian pregnancies were carrying fetuses with normal karyotype.

## Discussion

The prenatal diagnosis service in Estonia is funded by the National Health Insurance system and is thus accessible to all pregnant women, 99% of whom use this possibility. According to the latest Estonian national guidelines (https://www.ens.ee/ravijuhendid; available in Estonian), the standard protocol includes the combined testing based on I trimester serum screening and ultrasound examination. In case of the elevated risk (≥1:100) for fetal chromosomal trisomy, the confirmatory invasive testing is recommended. However, due to the quite significant FP rate of CT around 90% of invasive tests turn out to be unnecessary post-factum (18). NIPT, that is aimed to solve this problem, is currently offered as an out-of-pocket paid service in Estonia and the testing is being performed in foreign laboratories. The aim of the current study was to adjust previously described NIPT protocol (12) and our in-house developed NIPTmer analysis pipeline for detection of common fetal aneuploidies (11) in order to create a basis for introducing NIPT in the Estonian public health setting.

We analysed 447 samples including “high-risk” and normal pregnancies and correctly identified all non-mosaic samples aneuploid for chromosomes 21 (15/15), 18 (9/9) and 13 (4/4). In the current study z-score cut-off of 3 was applied for T21, T18 and T13. The relative coverage (or z-score) distribution among euploid and aneuploid pregnancies probably depends on both technological (cfDNA extraction method, sequencing library preparation, sequencing technology, and equipment used) and biological (gestational age, BMI and overall health of the mother) factors. Thus, the actual cut-offs for calling the elevated risk of a trisomy should be defined in every NIPT laboratory separately (11). These cut-offs determine the predictive power and the proportions of FP and false negative (FN) NIPT results. Setting cut-off value stringently would minimize the number of false-positive results and eventually enables to reduce significantly the number of invasive prenatal tests, which is one of the issues addressed by NIPT. However, one should keep in mind that this would introduce a number of false-negatives that must be avoided. With the current threshold value of “3”, six FP results (1.3%) were introduced. The patients in our study did not received any feedback on NIPT results. However, in case of a medical service the confirmatory invasive test would be offered to them (17) as NIPT represents a screening not a diagnostic test, though much more accurate than other fetal screening tests.

In addition to the common trisomies in our patients’ cohort, a case of mosaic trisomy 8 (47,XY,+8(11)/46,XY(29)) was detected by CVS but not by NIPT. Inability to pick up this abnormality can be explained by < 30% mosaicism and relatively low FF (6%), as well as the fact that in this study we only concentrated on three common trisomies. Generally, z-score cut-off determination for other chromosomal conditions is complicated as those cases are extremely rare. Although, the detection of chromosomal abnormalities other than T21, T18 and T13 (sex chromosome aneuploidies, other autosomal trisomies, triploidy, microdeletions) is included in some commercial NIPT platforms (e.g. MaterniT21™ by Sequenom and Panorama™ by Natera), ethical debates on whether this is of clinical utility are continuing. While proponents claim that in an era when broad genomic screening is possible there is no reasons to limit the testing to common trisomies, the opponents argue that with expanded cfDNA testing the current NIPT sensitivity of ~99% will decrease to ~60% leading to increased FP rate and therefore accompanied by additional challenges in counselling, increasing parental anxiety, unnecessary invasive testing, the potential for unnecessary terminations, additional costs, and decreased clinical relevance. Presently, the American College of Medical Genetics and Genomics (ACMG) discourages the use of NIPT to screen for autosomal aneuploidies other than those involving chromosomes 13, 18, and 21 (17). Although, routine cfDNA screening for microdeletion syndromes is not recommended by the American College of Obstetricians and Gynecologists (16), some authors find that testing for recurrent clinically relevant conditions (e.g. 22q11.2 deletion) may have larger clinical utility than traditional testing for T18 and T13 that are mostly lost in utero or soon after birth (19, 20). In our cohort, there were no pregnancies with rare autosomal trisomies (except for aforementioned mosaic trisomy 8) nor segmental imbalances, therefore we cannot estimate the readiness to detect such kind of abnormalities by our NIPT method and analysis pipeline.

With regard to sex chromosome aneuploidies (SCA), four cases of monosomy X were correctly determined with current NIPT pipeline, however, a case of 47, XXX, a syndrome with clinically mild phenotypic changes, was not detected. Generally, screening for SCA is also controversial. Some find that phenotypes associated with SCA do not qualify as severe diseases, and often infertility is the only problem. Testing for SCA for informational purposes only may therefore violate the right of future child to genetic and reproductive privacy, and additionally does not warrant public health care spending if the test is state funded (20, 21). Based on ethical and social considerations, fetal sex is not communicated to Dutch women undergoing NIPT testing, as there is no underlying health benefit. Information regarding sex chromosome aneuploidies also remains undisclosed because of biological difficulties, such as mosaicism, and clinical issues, such as less severe/or wide clinical consequences associated with these conditions (22, 23). Reiss et al. reported that getting to know that their fetus might have an SCA some patients regretted that they had learned this information prenatally, meaning that careful pre-test discussion explaining NIPT limitations is extremely important (24, 25). At the same time, in certain populations about half of the women carrying fetus with SCA detected by NIPT and confirmed by invasive test elect for termination of the pregnancy (26, 27). Basically, NIPT is less accurate in prediction of SCA compared to common trisomies. Some studies find that FP rate is especially high for Turner syndrome – up to 60-90%, which can be partially explained by maternal mosaicism for monosomy X which can be a part of normal aging (24, 28, 29).

Along with the issue to what extent one should be tested, another question exists, regarding the possible target groups for NIPT, which is also relevant for Estonia. It was demonstrated that there is no significant difference in NIPT screening performance between high-risk, routine and mixed populations (30). Therefore, the question arises what is the better way to implement NIPT into clinical practice and who are the main target groups for this testing.

In general, three models have been proposed for incorporation of NIPT into current screening programs (31). The first one assumes that NIPT is used as a second-tier screening test where women first undergo conventional screening and those with a high risk are subsequently offered NIPT or invasive testing. In this case, NIPT would be offered to ~5% of women, thus reducing costs (32). In the second model, the role of a universal primary screening test is reserved for NIPT in combination with ultrasound examination (33). This provides the highest detection rate of aneuploidy but at a higher cost. According to the third possible model, women deemed to be at high risk (e.g. ≥ 1:100) after conventional CT would be offered to choose between NIPT, invasive testing or no follow-up, and those with the intermediate risk (e.g. 1:101-2500) would be offered NIPT or no further testing. This model would be a balance between cost and detection rate (34).

Various national studies evaluating NIPT performance demonstrated that NIPT enables to decrease significantly the number of invasive testing. For instance, according to the results of TRIDENT study conducted by Dutch NIPT consortium, the reduction in invasive testing was at least 60% (22). In general, depending on the country and the screening population, the reduction in invasive testing after introduction of NIPT achieves up to 75% (35). Thus, NIPT fulfills its primary goal, namely the reduction in the number of invasive test procedures. Although, there is still controversy on the magnitude of the fetal loss rate caused by invasive testing, the reduction of invasive procedures is certainly perceived as better care by both health care providers and pregnant women (22). Furthermore, the implementation of NIPT decreases the median gestational age at time of diagnosis and therefore the gestational age at time of abortion for aneuploidy. In addition, with introduction of NIPT the number of high-risk pregnant women refusing further testing also drops (36). Taken together, these studies support introduction of NIPT as a safe second-tier screening test to select accurately the small proportion of women truly at high risk for fetal trisomy. A prerequisite here is existing publicly available first-trimester CT for the primary selection in a country of interest. Amongst others, this requirement is fulfilled in Estonia.

However, the use of NIPT as a secondary screening test mainly lowers the FP rate, while does not diminish the FN rate of the initial CT. This could be overcome by replacing conventional serum analysis by NIPT as a first-tier screening test combined with ultrasound examination (second model) due to the higher detection rate of NIPT compared to conventional CT (22). Recently, this turned an option of choice in a number of countries, including The Netherlands and Belgium where NIPT for the common trisomies is offered to all pregnant women since 2017. Regardless of which NIPT implementation model is chosen, every positive NIPT result must be confirmed using invasive testing before any further actions are undertaken, because in all cases NIPT represents a screening tool not a diagnostic test.

In the current study, novel in-house developed NIPTmer program was implemented for fetal aneuploidy detection. NIPTmer is about an order of magnitude faster than mapping-based NIPT tools, uses less computer resources and does not require previous experience with mapping NGS reads. It allows on-the-fly updating of both control and aneuploidy datasets so the prediction accuracy gets the better the more analyses are performed. The program is also more robust as the non-unique and polymorphous parts of genome are discarded before the actual analysis, in k-mer list generation step, and thus do not introduce additional variability in test results.

Reference group for NIPTmer analysis (“general population” group) in our study consisted of patients with wide range of gestational age (10 to 40 weeks). No limitations on gestational age were implemented to reference group in order to validate NIPTmer and SeqFF programs in various conditions including low and high FF. “High risk” group, however, were comprised of patients with gestational age suitable for NIPT analysis prior to invasive testing (11-21 weeks).

In conclusion, we validated next-generation sequencing based NIPT method and original NIPTmer software program among Estonian population. Our NIPT methodology proved to perform efficiently in detecting common fetal aneuploidies T21, T18 and T13 with high sensitivity and specificity even in case of low FF. Although improvements to the analysis pipeline could be added to identify rare aneuploidies, the test is ready for implementing into the prenatal clinical care in Estonia.

## Acknowledgements

The authors are very grateful to the patients who participated in this study.

